# A principled approach to the identification of biologically relevant transcription factor binding sites

**DOI:** 10.1101/2025.09.25.678593

**Authors:** Maalavika Pillai, Luís A. Nunes Amaral

## Abstract

Transcription factors (TF) regulate gene expression in cells by binding to specific DNA sequences. However, previous studies indicate that TF binding is a necessary but not sufficient condition for regulating gene expression.^1–3^ While recent advances in sequencing technologies such as Chromatin Immunoprecipitation sequencing (ChIP-seq) enable researchers to identify binding sequences at single nucleotide resolution, they often fail to distinguish between biologically relevant and non-relevant – i.e., spurious – binding events. Reliably distinguishing between these two types of binding is essential if one aims for reproducible findings. Here, we report on a systematic study of the sensitivity of “called” TF–target gene interactions to the choice of various data processing parameters. We find that acceptable distance from the transcription start site (TSS) is a critical parameter for reducing subjectivity. We, thus, develop a two-state mechanistic model that captures the positional distribution of both biologically relevant and spurious binding events, allowing us to identify interaction distance thresholds that minimize spurious binding. We validate our model with independent data by showing that biologically relevant binding events identified by our model are capable of recruiting RNA Polymerase II and, subsequently, regulating mRNA expression levels of the target gene. Finally, we investigate the impact of our approach on the detection of genes that are differentially bound and regulated by BCL3, a transcription coactivator, during molecular perturbation experiments. Within the genes expected to show differential expression, our model predicts that over 70% of reported interactions are spurious and we demonstrate the lack of change in mRNA expression in these genes following perturbation. We also identify genes that display a shift from spurious to biologically relevant binding that account for close to 17% of the genes displaying a sharp change in mRNA expression and strong association with known functions of BCL3 such as cell cycle regulation and DNA damage response. By providing a systematic TF-specific method to identify binding events that are directly involved in transcriptional regulation our model will increase reproducibility of analysis.

## 1 Introduction

Transcription factors (TFs) are DNA binding proteins that bind to specific sequence motifs near the transcription start site to regulate the transcription of their target genes. One of the most common methods to identify these binding sequences motifs is Chromatin Immunoprecipitation Sequencing (ChIP-seq) which uses antibodies against a specific TF to isolate it in DNA-bound form. The DNA bound to the TF is then sequenced and aligned to a reference genome to identify TF binding sites at single nucleotide resolution and uncover target genes. ChIP-seq data is used to identify recurring motifs or enriched binding sequences for specific classes of TFs.^4,5^ The TF–target gene interactions identified from ChIP-seq data can also be aggregated to construct the gene regulatory networks (GRNs) that control the global expression of various genes within a cell.^6,7^ ChIP-seq is also often used in molecular perturbation experiments to identify differential binding events that regulate gene expression.^8,9^

While ChIP-seq data has the potential to help ascertain the biological role of TFs in a cell, previous studies suggest that many computational pipelines identify large numbers of so-called “phantom peaks” — false positive signals.^10^ Phantom peaks can occur in genomic regions that are interaction prone or “sticky.” These sticky regions frequently display binding events from multiple TFs.^11^ Phantom peaks can also occur due to weak or transient TF binding that does not lead to active transcriptional regulation but nonetheless gives rise to a ChIP-seq signal.^12–14^ Phantom peaks can result in a lack of reproducibility across multiple replicates of the same experiment.^15^ Thus, there is a clear need for adequate processing of ChIP-seq data to ensure that downstream analysis can be used to make accurate inferences.

Several tools and resources have been developed for helping to identify recurrent motifs or enriched binding sequences from ChIP-seq data. *Homer* and MEME-ChIP^16–18^ are commonly used tools for identifying consensus binding sequences or probability weight matrices that determine the probability of binding to each nucleotide at a specific position. JASPAR and TRANSFAC are resources that provide access to probability weight matrices for a large number of TFs. The Encyclopedia of DNA elements (ENCODE), NCBI Gene Expression Omnibus, EMBL’s European Bioinformatics Institute are publicly accessible databases holding large numbers of experimental datasets. These resources often use uniform processing pipelines to ensure reproducibility and minimize experimental bias, making them particularly helpful for validation and benchmarking. For instance, JASPAR uses fixed probability weight matrices (PWM) score and p-value thresholds — PWM ≥ 0.8, *p <* 0.05 — when predicting transcription factor binding sites across 8 organisms.^4^ Databases such as ENCODE follow strict data processing pipelines that not only require all sequencing data to be processed uniformly but also mandate experimental standards pertaining to reproducible replicates, appropriate controls and sequencing coverage.

Other forms of data-processing standards are often integrated into signal detection or peak-calling tools that distinguish signal from noise in sequencing data. For example, *Macs2* integrates a statistical model that accounts for the difference between tag density on the Watson and Crick strands to identify regions that have enriched tag density.^19,20^ Another method that is commonly used to remove problematic regions in the genome is by “blacklisting” or annotating these regions and discounting signals in these regions. To this extent, ENCODE utilizes a list of curated blacklisted regions for their datasets which can also be defined during peak calling with programs such as *Macs2* for independent ChIP-seq experiments.^21^ However, this method requires prior knowledge of these problematic regions which might not be possible for datasets arising from under-studied species or underrepresented contexts. Unfortunately, even datasets such as ENCODE, that integrate these bias correction methods, still have not resolved^10,15^ the lack of reproducibility challenge, demonstrating the need for additional work.

Here, we report on a principled approach to the detection of bindings events in ChIP-seq data. Our modeling assumes two mechanisms for the presence of TF binding domains in the genomic regions surrounding the transcription starting site (TSS) of genes – the first process is just random occurrence through genome drift, the second is selection for domains that occur close to the TSS of target genes whose regulation by the TF provides a fitness advantage. Our model yields a straightforward, statistically-sound way to decide whether a called peak is biologically-relevant or spurious.^1^ We independently confirm the validity of our classification of TF–target gene interactions by considering recruitment of RNA polymerase II to the TSS of the target gene and by evaluating correlations between mRNA abundances of the TF and the target gene. Finally, we demonstrate that in A549 cells treated with 100nM dexamethasone, close to 70% of differential binding events for transcription factor BCL3 fail to elicit a change in mRNA expression and these interactions are accurately captured as spurious binding events by our model.

## 2 Modeling

### 2.1 Sources of variability

The sources of variability in the identification of biologically relevant TF–gene interactions from ChIP-seq data can be largely classified into three classes based on the stage of sample processing (**Fig. 1**). The first arises from intrinsic variation in TF binding sites across various samples and other fluctuations that accumulate during the immunoprecipitation stage. These include differences in TF binding due to single nucleotide variant allele specificity or asymmetric binding across strands leading to lack of concordance across replicates^3,23,24^, differences in crosslinking and fragmentation efficiency^25–28^ and differences in antibody specificity towards isoforms/mutants or lack of TF-antibody titration to account for variable TF abundance.^29–31^ The second arises during DNA sequencing and include low read depth, poor sequencing coverage, PCR duplication, poor alignment of repeat sequences and non-unique alignment.^27,32–35^

**Figure 1.**
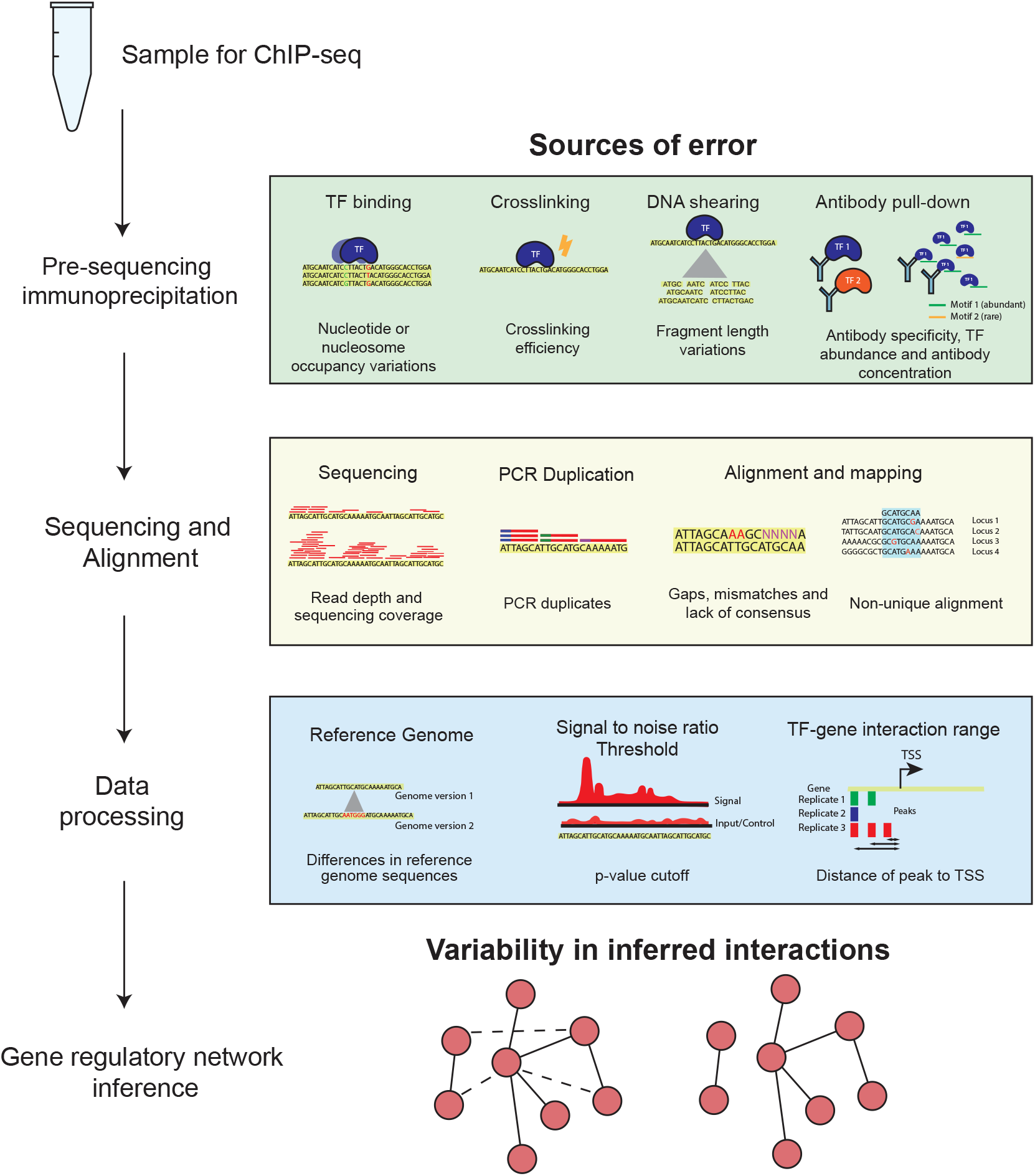
ChIP-sequencing data has multiple sources of error that can lead to spurious TF-gene interaction detection. Error sources during ChIP-seq can be classified into three groups based on the stage of sample processing: pre-sequencing immunoprecipitation, sequencing and alignment, and the sequencing data processing. In this paper, we focus on the variables affecting the last stage, sequencing data processing, which includes sequence variations between reference genome versions, the signal-to-noise threshold for peak calling and the distance between the TF binding site and the TSS of the target gene to define an interaction.

Lastly, and of interest to us here, the third arises from computational pipeline choices including differences in alignment results due to use of different reference genomes^36,37^, in the choice of an appropriate signal-to-noise ratio threshold^19,38^, and in the cutoff for determining sufficient physical proximity to the transcription starting site (TSS).^39^ For example, selection of uniform data processing parameters across datasets may fail to account for differences in peak or signal abundances across experiments. These errors can further percolate into downstream analysis and lead to poor reproducibility and inaccurate inferences (**Fig. 1**).

To further understand the effects of data processing parameter variation on inferred TF-gene interactions, we conducted a systematic sensitivity analysis for ChIP-seq data for 5 cell types observed during embryonic haematopoietic differentiation.^40^ We then developed a two-state statistical model to determine appropriate interaction distance thresholds that filter out spurious binding events, i.e., binding events that are not associated with transcriptional regulation. Our model identifies dataset-specific parameters for distinguishing biologically-relevant from spurious binding events that hold up against a wide range of ChIP-seq datasets with varying degrees of signal abundance. Importantly, our model enables researchers to make decisions based on setting a false detection rate (FDR) that is consistent with the goals of the downstream analysis. Lastly, we validate our model against other modalities of gene expression such as RNA Polymerase II co-localization and RNA-sequencing. These validations confirm the ability of our model to accurately identify transcriptionally relevant peaks in ChIP-seq data.

### 2.2 The spatial distribution of TF binding sites

Let us reason about how the position of a potential binding site for a specific TF around a specific gene may evolve over time. For concreteness, let us assume that there is a set of *k* short sequences 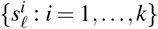 of length *ℓ* base pairs that bind effectively with the TF under consideration. Now, consider the DNA sequence *S*_*g,L*_ extending for *L* base pairs in both directions from the transcription starting site (TSS) of gene *g*. Over evolutionary time scales, this sequence will be changed through mutations, insertions, deletions, and crossovers.

If there is no selection pressure for the presence or absence of any of the 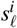, then the mechanisms changing the genome will contrive so that any emerging sequence performs a random walk within *S*_*g,L*_. After a long enough time, such diffusion will give rise to an effectively uniform spatial distribution of the 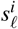. That is

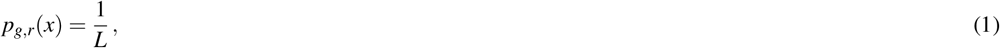

where the *r* refers to random diffusion, *x* ∈ {0, …, *L*} is the distance to the TSS of a sequence in the set 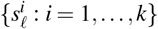 and there is no preference in the diffusion for the upstream or downstream directions.

Let us now assume that there are some genes *g* that are required to be regulated by the specific TF considered here. Then, there will be selection pressure for having binding sites physically close to the TSS. That is, diffusion of potential binding sites will no longer be random, but there will be instead a bias toward the TSS. If we model this bias as a potential well, then the diffusion process yields an exponential spatial distribution of the binding sequences

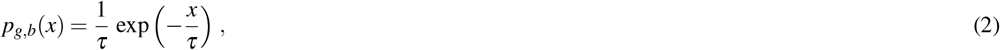

where the *b* refers to biased diffusion, *x* ∈ {0, …, *L*}, and *τ* ≪ *L* is related the magnitude of the selection pressure for the regulation of the gene by the TF.

To put these together, we will assume that there is some fraction *f* of genes that are regulated by the TF. Under the assumption that every gene experiences the average selection pressure for regulation by this TF, the overall spatial distribution of bindings sites at distance *x* from the TSS is given by

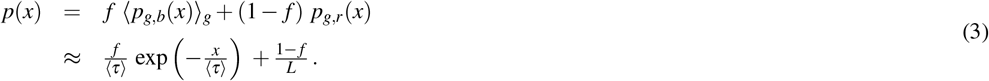

We acknowledge that the underlying assumption used here is unlikely to be strictly true. Unfortunately, this is also an assumption that cannot be resolved experimentally with available data, so we will drop the ⟨… ⟩ from our notation. Importantly, if this assumption is approximately correct, then one would expect that *τ* would take the same value for all equally important TFs.

For comparison with empirical data, it is best to consider not Eq. (3) but the cumulative distribution *P*(≤ *x*). Moreover, since the number *N* of detected binding sites will depend on the abundance of the TF, we focus on this number instead of the distribution

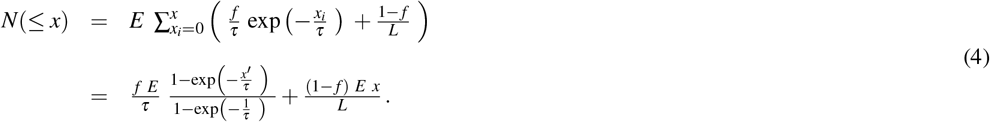

The parameter *τ* can be easily estimated from the data as can 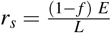, which quantifies the spurious binding rate, and 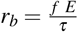, which is the scaling factor for the biologically relevant binding rate (**Fig. 2a-b**).

**Figure 2.**
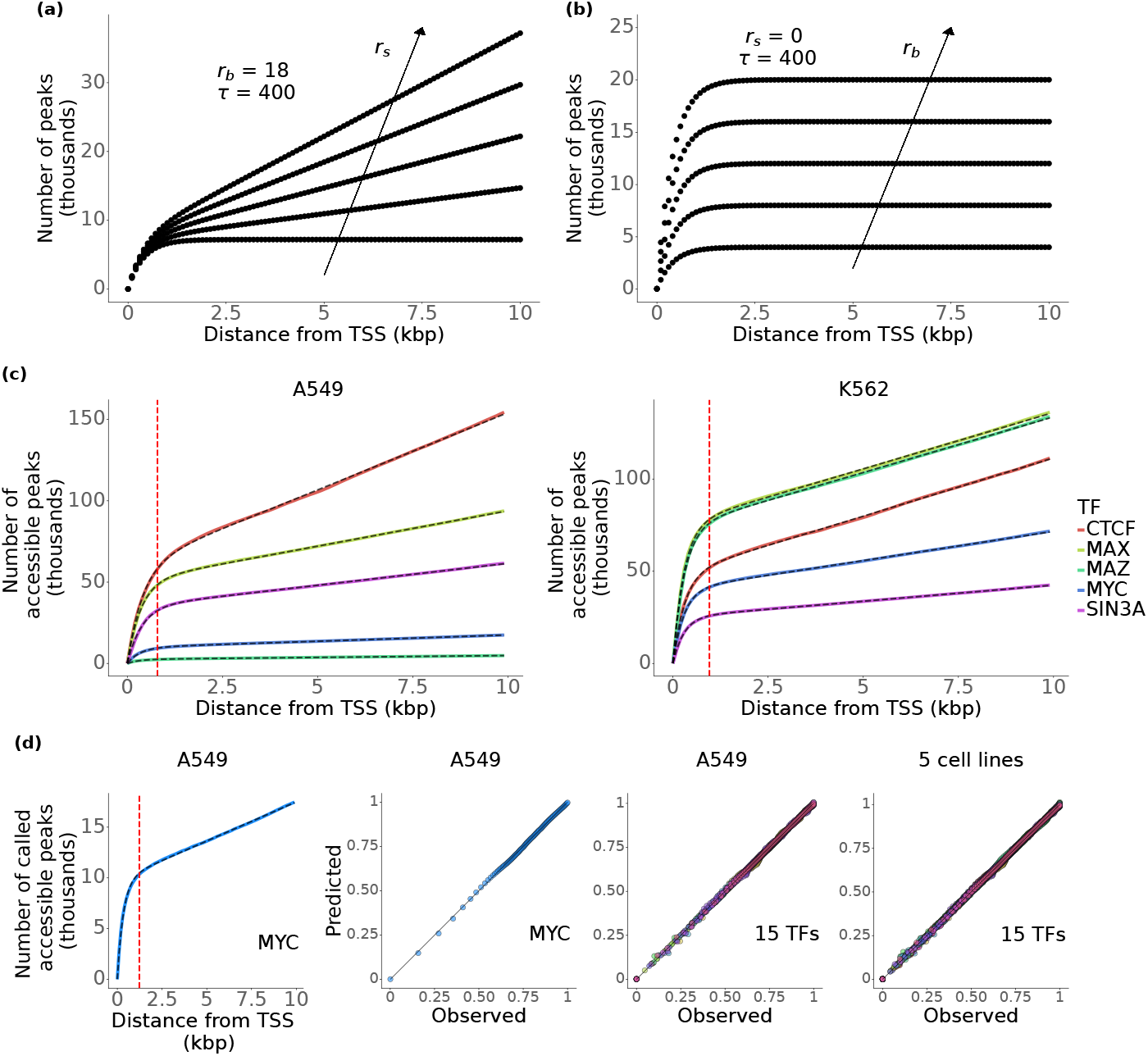
Exponential decay with constant spurious binding accurately captures the cumulative distribution of called peaks with respect to varying interaction distance thresholds. **(a)** Increasing the spurious binding rate (*r*_*s*_) increases the slope of the linear phase of the cumulative distribution for a fixed number of true targets (*r*_*b*_ and *τ* are constant). **(b)** Increasing the scaling factor of biological binding rate (*r*_*b*_) increases the total number of targets that a TF binds to in the absence of spurious binding (*r*_*s*_=0) and with a fixed decay rate (*τ*). The spurious binding rate (*r*_*s*_) changes the slope of the second phase of the curve while *r*_*b*_ controls the total number of true targets a TF can bind to. **(c)** Cumulative number of called peaks for varying distance from TSS displays two distinct phases across TFs (different colors) and cell lines A549 (left) and K562 (right). Close to the TSS, in the first phase, the curve follows a cumulative exponential distribution. At large distances from the TSS, the second phase, the cumulative number of called peaks is linear indicating a constant rate of spurious binding. Red dashed line indicates average value of *x*_*o*_ across all TFs plotted for each cell line. Dashed black line represents the fit for each curve. **(d)** Comparison between model fit and observed number of called peaks. For each TF ChIP-seq dataset, the number of called peaks is normalized by the total number of observed peaks to allow for comparisons across multiple datasets. The model fits closely follow the observed number of called peaks for all 15 TFs across the 5 cell line.

## 3 Data and Methods

### 3.1 ChIP-seq data analysis

For exploring the dependence of ChIP-seq data analysis on different data processing choices, we retrieved publicly available GEO dataset GSE69101^40^ for downstream analysis. The data comprises ChIP-seq data for 16 TFs across 5 hematopoietic developmental stages – Hemangioblasts (HB), Hemogenic endothelium (HE), hematopoietic progenitors (HP), Macrophages (MAC) and Mesoderm (MES). Specifically, we downloaded *fastq* files and aligned reads against two mouse genome releases: GRCm38 (mm10, GenBank assembly GCA_000001635.2) and GRCm39 (mm39, GenBank assembly GCA_000001635.9). We identified binding peaks using the *callpeak* function in *Macs2*.^19^ We considered several p-value thresholds in order to select a desired signal-to-noise value. For most analysis, we settled on *p* = 10^−5^.

For evaluation of our model for the spatial distribution of TF binding sites, we retrieved ChIP-seq data from ENCODE.^41^ Specifically, we downloaded ChIP-seq datasets for 15 TFs (Table 1) across 5 cell lines (A549,HepG2, GM12878, K562 and MCF-7).

For A549 cells treated with 100nM dexamethasone, we downloaded ChIP-seq datasets for 7 TFs that were available for both treated and untreated samples. While the analysis shown in Figure 5 pertains to a single transcription factor (BCL3), binding events from all 7 TFs were used to identify the control group (Not Expected/Not Predicted genes) to avoid capturing any changes associated with the activity of other TFs.

To estimate the number of accessible binding sites, we measured the overlap in the ChIP-seq peaks and the corresponding ATAC-seq dataset for each cell line using *bedtools*. Only accessible peaks within 10 kbp of the TSS were retained for further analysis. The absolute distance from the center of each peak to the TSS of the target gene was used to estimate the interaction distance. The list of target genes was obtained from the reference genome used for alignment. For each dataset, the number of edges was estimated as the number of unique genes that a TF interacts with.

We define the interaction distance threshold *x*_*o*_ as the distance at which the ratio of biologically relevant binding to spurious binding events is 10.

We classified the interaction between a TF and a target gene to be biologically-relevant if we identify at least one TF binding event — that is, a peak in ChIP-seq data — within the interaction distance threshold from the gene’s TSS, otherwise, the interaction was deemed spurious.

### 3.2 RNA-seq data analysis

To test the influence of a transcription factor on the expression of its target genes, we measured pairwise correlation in mRNA expression for each pair of TF and target gene. For this analysis, we only considered K562 since it was the only cell line with more than 30 samples.

We obtained estimates for mRNA expression using RNA-sequencing data for K562 cell line from ENCODE^41^ and estimated Spearman correlation coefficients for each TF-target gene pair. We determined the correlation for a TF-target gene pair to be significantly and strongly correlated if the absolute value of correlation coefficient was greater than 0.5 and the p-value was below 0.05.

For A549 cells treated with 100nM dexamethasone, we downloaded RNA-seq datasets from ENCODE for treated and untreated samples from the same experiment. The average log2 fold change in mRNA counts between treated and untreated sampleswas estimated for each gene in each class.

### 3.3 RNA Polymerase II co-localization

To test if a TF binding event could recruit RNA Polymerase II and initiate transcription, we measured RNA Polymerase II localization at the TSS of all target genes of a TF. We estimated RNA Polymerase II recruitment using POLR2A (RNA Polymerase II Subunit A) ChIP-seq data from ENCODE for each of the 5 cell lines.^41^ We deemed that there was RNA polymerase co-localization if there was a POLR2A peak within TSS +/-2kbp of all genes containing a TF peak.

We retrieved POLR2A bigwig files from ENCODE in order to visualize RNA Polymerase occupancy (**Fig. 3**). We aggregated signal intensity within bins of size 10 bp for the region within 2 kbp of the TSS using *deepTools*.^42^ For each target gene, we then created a heatmap of the scaled signal scores of the RNA Polymerase II occupancy as a function of the distance to the TSS.

**Figure 3.**
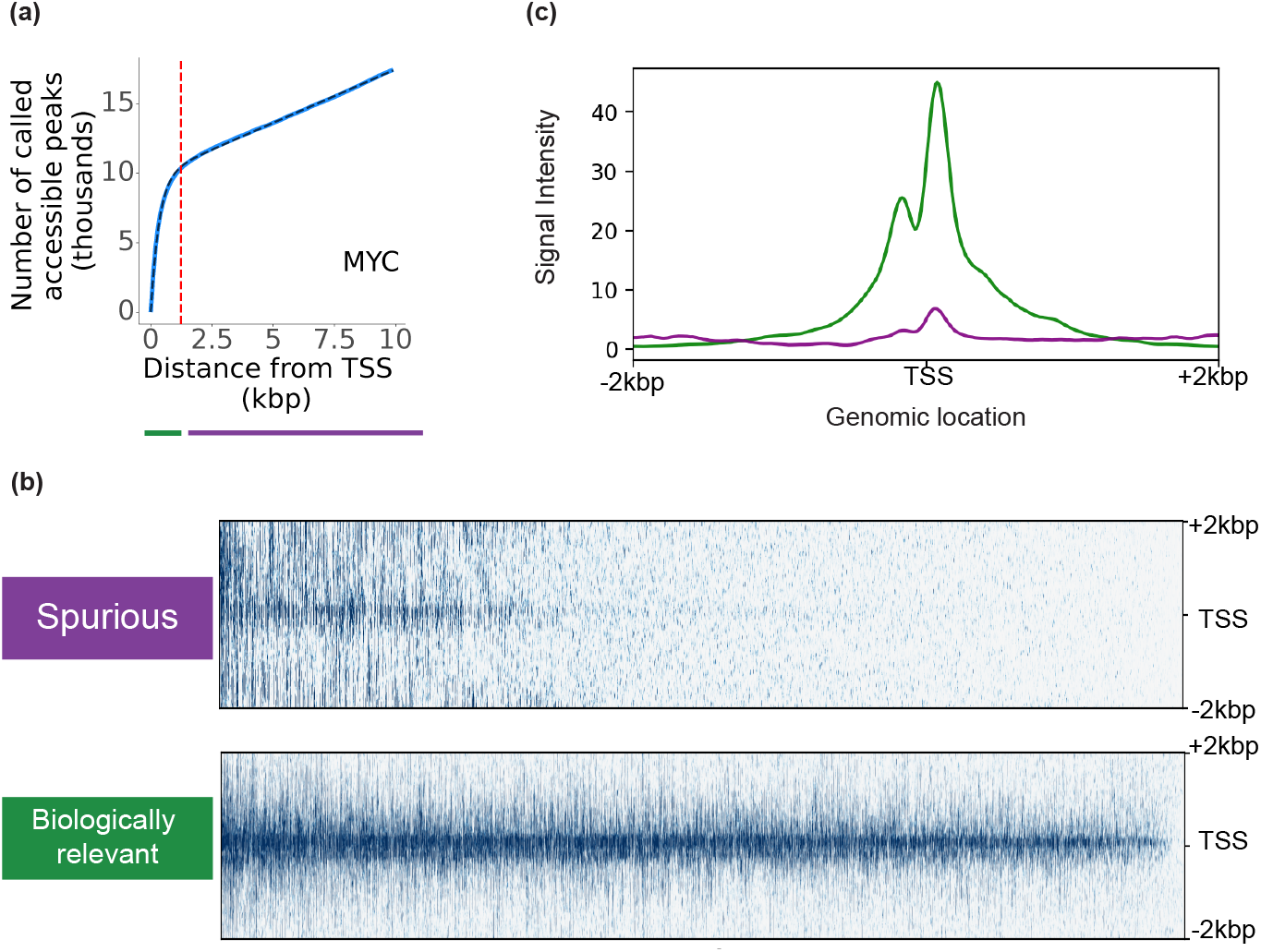
Biologically relevant peaks identified by the model display RNA Polymerase II recruitment. **(a)** A classification threshold was estimated for each dataset. **(b)** RNA Polymerase II displays strong occupancy signal at TSS of target genes that have a true peak based on the model. In contrast, target genes with only model-based spurious peaks show weak to no RNA Polymerase II recruitment at TSS. **(c)** Distribution of RNA Polymerase II ChIP-seq signal intensity for target genes with true peaks (green) and spurious peaks (purple).

### 3.4 Functional pathway enrichment analysis

To test for enrichment of biological pathways in expected and predicted target genes of BCL3, we conducted Fisher’s exact test against all Gene Ontology (GO) terms associated with biological processes using *gProfiler*.^43^ We used g:SCS multiple testing correction method to account for dependence between GO terms and a significance threshold of 0.05 to identify hit terms.

## 4 Results

### 4.1 Signal-to-noise threshold and interaction distance range determine number of TF-Gene interactions inferred from ChIP-seq data

Inferring TF-Gene interactions from ChIP-seq data involves choosing appropriate data processing parameters that capture biologically relevant interactions and minimize noise. To measure the effect of these choices on the inferred gene regulation networks (GRNs), we tested the sensitivity of the number of identified interactions to changes in the reference genome version, the signal-to-noise threshold, and the interaction distance range. As detailed in the data and methods section, we analyzed the publicly available GSE69101 dataset and compared the network inferred from our analysis to that reported in the original study.^40^ For estimating the sensitivity to changes in each of the processing pipeline hyper-parameters, we obtained the total number of peaks called and the number of unique TF–gene interactions (GRN edges) obtained by the two pipelines.

#### Reference genome

We compared the number of peaks and GRN edges for reads aligned to two mouse reference genomes: GRCm38 (mm10) and GRCm39 (mm39). As hoped for, we observe no significant difference in either of these metrics upon changing the reference genome version (**Fig. S1**).

#### Signal-to-noise threshold

We compared the number of peaks and GRN edges when changing the ‘*-p*’ argument in the *callpeak* function in *Macs2*.^19^ We considered values from 10^−9^ to 10^−1^, a range covering the values previously reported in literature.^40^ As expected, we observe a significant and exponential increase in the number of peaks and the number of GRN edges when setting higher — that is, more lenient — *p*-value thresholds (**Fig. S2**).

#### Threshold for interaction distance

We compared the number of peaks and GRN edges when changing the interaction distance from the TSS for considering a binding event to be involved in transcriptional regulation of a gene. We considered values from 0 to 10 kbp. As expected, we see an increase in the number of peaks and edges that are identified as the threshold interaction distance increases (**Fig. S3**).

#### Impact of sensitivity

Consistent with expectations, we thus find that the number of TF-Gene interactions identified can change drastically with choice of *p*-value and threshold for interaction distance. Such extreme sensitivity raises concerns about how one ensures reproducibility of reported results and suggests the need for the adoption of a standard pipeline and appropriate reporting of hyper-parameter values.

To explore this concern, we next compared the network inferred in the original paper to that inferred from our analysis. We identified TF-Gene interactions that were both reported by Goode *et al*. and identified with our analysis using the signal-to-noise threshold reported by Goode *et al*. and a threshold interaction covering the coding sequences of the 16 target genes discussed in the paper. Note that the value of this hyper-parameter was not reported by Goode *et al*. Surprisingly, while there were only two edges identified in our analysis but not reported by Goode *et al*., there were close to 30 edges reported by Goode *et al*. but that were not identified in ours (**Fig. S4a, Table 2**). Remarkably, for many of these 30 edges, the identified TF peak was farther than 50 kbp from the TSS (**Fig. S4b, Table 3**). While one could hypothesize that such peaks might indicate the presence of distal cis-regulatory elements, it is worth reporting that some of those distant peaks that were reported as interactions had been previously reported in other publications by some of the authors of Goode *et al*.

### 4.2 Two-state model captures positional distribution of transcription factor binding

We use publicly-available ChIP-seq data from ENCODE for five cell lines (A549, HepG2, GM12878, K562, MCF-7) and 15 TFs.^6,41^ We then measure the cumulative number of TF binding events as the number of peaks that are “called” up to a given distance from the TSS and find results consistent with the two-state model we described earlier (**Fig. 2c, Fig. S5**). Specifically, we find that while the results for different TFs within the same cell line have the same functional form — rapid increase followed by linear growth^39,44^ — they can differ quite dramatically in the parameter values. The same patterns is also visually apparent when considering the same TF but across different cell lines, suggesting differences in target genes across different tissue-specific and environmental contexts.

We next fit the two-state model to each TF–cell line pair in the ChIP-seq datasets. The initial rapid increase representing the contribution of the biologically-relevant exponential decay term, characterized by the parameters *r*_*b*_ (biologically-relevant binding rate) and *τ* (the decay constant), and the linear growth regime corresponding to the uniform spurious binding, characterized by the parameter *r*_*s*_ (spurious binding rate). The biologically-relevant binding rate reflects the total number of “true” target peaks for the corresponding TF. Higher spurious binding rate, *r*_*s*_, increases the slope in the linear regime (**Fig. 2a**). Higher *r*_*b*_ yields a higher number of true target peaks (**Fig. 2b**). For visual clarity we do not plot the 15 × 5 curves for all TF–cell line pairs. Instead, we normalize by the maximum observed value both the model predicted and the observed cumulative number *N*(*> x*)?of called peaks. In Fig. 2d, we plot first the raw values of *N*(*> x*)?for MYC in cell line A549. Next, we plot the normalized values, demonstrating the close agreement between model and data for this TF in this cell line. We can then add to the plot the results for all other 14 TFs in A549 and, finally, all 15 TFs for all 5 cell lines. The close agreement across these 75 cases suggests that our two-state model accurately captures the patterns in the data.

#### Characteristics of estimated model parameters

We calculate the distributions of estimated model parameters from the 75 TF–cell line pairs. The width of these distributions suggests TF and cell line dependence. To further investigate such dependence, we also visualized the range of values each parameter takes for a given TF across the 5 cell lines (**Fig. S6-8**). We find that the biologically-relevant binding rate, *r*_*b*_ (**Fig. S6**), and the spurious binding rate, *r*_*s*_ (**Fig. S7**), can vary by 10-fold across TFs, but have much smaller variability when looking across cell lines for the same TF. In contrast, the decay rate, *τ* (**Fig. S8**), varies by less than 2-fold across TFs, and even less when looking across cell lines for the same TF.

Biologically, the variation across cell lines in the estimated values of *r*_*b*_ and *r*_*s*_ for a single TF can be rationalized as the consequence of differences in accessibility of target binding sites that might exist across cell lines and expression levels of the TF. For instance, a cell line with accessible regions containing higher instances of a TF’s binding motif and target genes is likely to have higher values of these parameters. We know that chromatin accessibility greatly differs for cell lines originating from different tissues and for different biological contexts (for instance, cancer vs. non-cancer samples).

The greater uniformity in the estimated values of the the decay rate, *τ*, also conforms to our expectation. Whereas changes in chromatin accessibility and expression levels of the TF may change the number of called peaks, the positional distribution of called peaks around the TSS will not be affected by those mechanisms. Instead, the only mechanism by which the length scale of the distribution would be changed is sequence changes in the genome — which will occur across samples from different species or on much longer time scales. ^45,46^ Such changes would not to be captured in the data we consider.

As a robustness test of these findings, we investigated how a change in the p-value hyper-parameter might impact the estimation of the two-state model parameters (**Fig. S9**). As expected, we observe large changes in the estimated values of *r*_*b*_ and *r*_*s*_, but not of *τ*, further strengthening our confidence in these findings and the suitability of the assumptions underpinning the two-state model.

### 4.3 Model-identified biologically-relevant interactions have greater RNA Polymerase II recruitment

We next further test the biological relevance of the predictions of our model using independent data. If the called peaks we classify as biologically-relevant or spurious (**Fig. 3a**) are indeed so, then we would expect then to display different rates of recruitment of RNA Polymerase II. To test this hypothesis, we measured RNA Polymerase II occupancy as the ChIP-seq signal for POLR2A– RNA Polymerase II Subunit A– in A549 around the TSS for each target gene of MYC. We then compared occupancy rates for biologically-relevant TF–target gene against binding rates for spurious TF–target gene interactions. It is visually apparent that there is a strong, statistically significant signal for identified biologically-relevant interactions, while for spurious interactions there is only a background signal (**Fig. 3b-c**).

### 4.4 Model-identified biologically-relevant interactions display higher correlations in the levels of TF and target gene mRNAs

If the called peaks we classify as biologically-relevant or spurious (**Fig. 3a**) are indeed so, then we would expect strong correlations between mRNA abundance levels of the TF and the target gene. In order to test this hypothesis, we measure the correlation between mRNA expression of a TF and its target gene using Spearman’s correlation coefficient and estimate the fraction of target genes displaying strong statistically significant correlations, which we define as a correlation coefficient with magnitude greater than 0.5 and *p <* 0.05. We find an unambiguous difference in the fractions of strongly correlated pairs (**Fig. 4a**). Whereas the correlation is strong and significant for over 50% of TF–target gene interactions classified as biologically-relevant, the fraction is smaller than 30% for interactions classified as spurious. The value of Spearman’s correlation coefficient is also higher in interactions derived from biologically relevant binding events (**Fig. 4b**), further suggesting that the model derived biologically relevant peaks accurately identifies transcriptionally co-regulated TF-target gene interactions.

**Figure 4.**
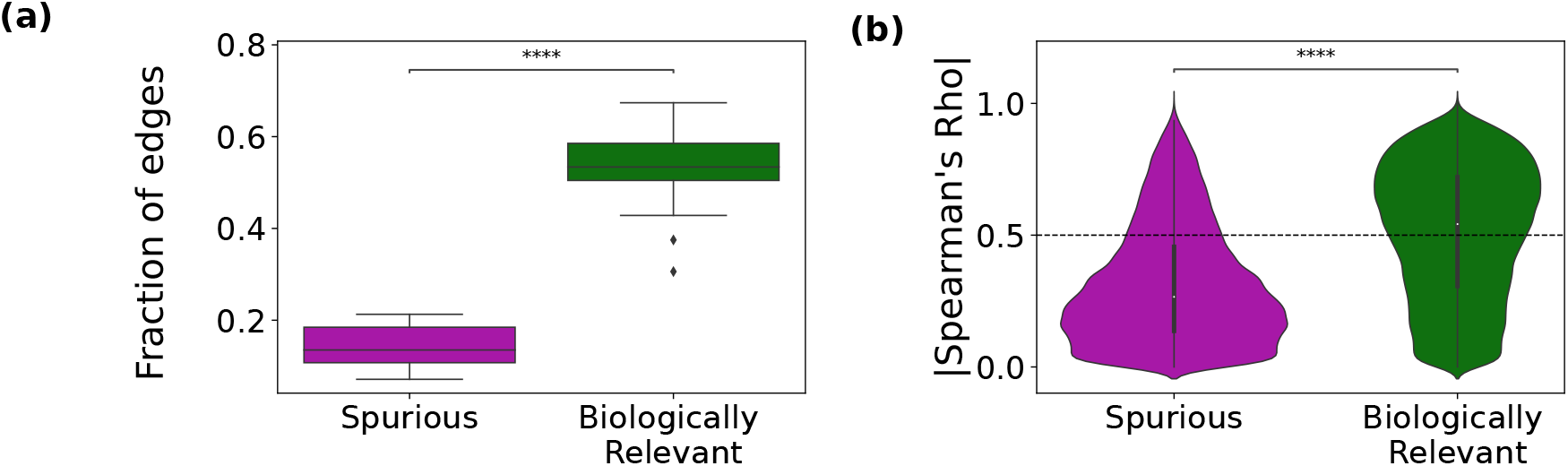
TF-Gene interaction pairs identified from model-derived biologically relevant peaks are transcriptionally correlated. (a) Fraction of TF-Gene interactions derived from peaks within(green) and beyond (purple) the interaction distance threshold (*x*_*o*_) that displayed statistically strong significant correlation ( |*ρ*| *>* 0.5, *p* ≤ 0.05) in K562 cell line RNA-seq data. (b) Distribution of absolute value of pairwise correlation coefficient values (Spearman’s rho) between a TF and its target genes for TF-Gene interactions derived from peaks within(green) and beyond (purple) the interaction distance threshold. The edges detected below the classification threshold (predicted true edges) display higher levels of RNA co-expression than the remaining edges (predicted false). **** indicates *p* ≤ 10^−4^

### 4.5 Two-state model predicts differentially expressed genes in molecular perturbation experiments

To test the ability of the model to identify true transcriptional targets of a TF, we compare the mRNA expression of genes expected to be differentially bound during a molecular perturbation with and without the model. Previous studies have reported extensive changes in gene expression and chromatin organization for A549 cells treated with 100nM dexamethasone (DXM)^47^. We used this molecular perturbation experiment to test if our model could accurately identify target genes that display changes in TF binding and subsequent changes in mRNA expression. We compared binding events for BCL3, CTCF and JUN in untreated control samples and 100nM dexamethasone treated samples. Each gene with a binding event within 10kbp of the TSS was classified as either biologically relevant or spurious based on the two-state model (**Methods**). Genes changing from no binding or spurious binding to biologically relevant binding from control to DXM treated (and vice versa) were predicted to be differentially expressed while genes shifting from no binding to spurious binding from control to DXM treated (and vice versa) were predicted to not have any change in expression levels. Without the two-state model, genes changing from no binding to binding (either biologically relevant or spurious) are expected to give rise to a change in expression levels (**Fig. 5a**). We also used a group of genes that exhibited no binding event within 10kbp of the TSS for any TF across both conditions as the control group (not expected and not predicted to change). Differentially bound genes identified without the model are classified as “Expected” or “Not Expected” while those identified based on the two-state model are classified as “Predicted” and “Not Predicted”.

**Figure 5.**
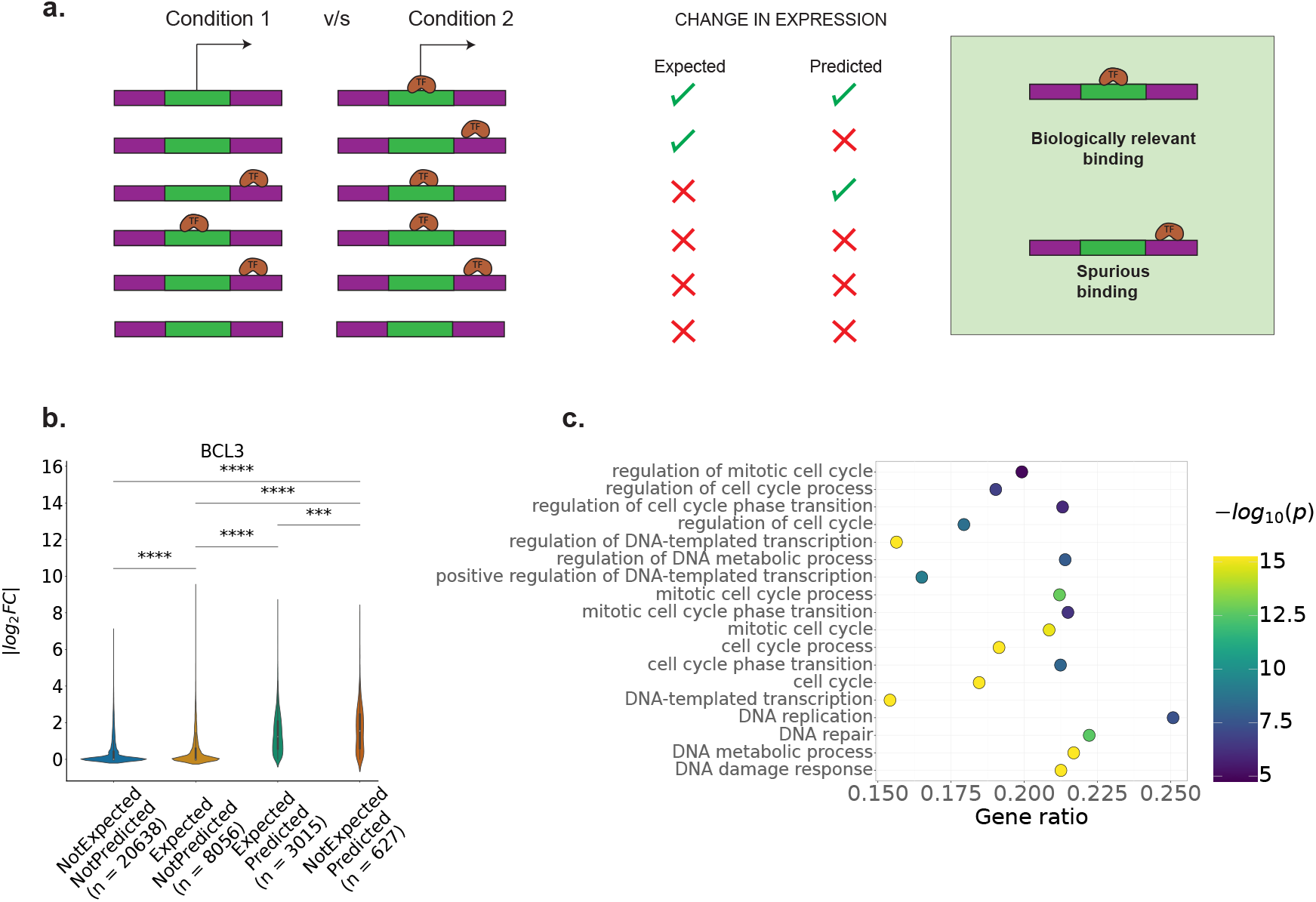
Two-state model captures differentially bound genes during molecular perturbation. A549 cells treated with 100nM dexamethasone for 12h display changes in TF binding events with and without treatment. **(a)** Schematic of genes expected to have differential binding without the two-state model v/s genes predicted to change with the model. **(b)** Distribution of average *log*_2_ fold change in mRNA expression between treated and untreated samples. Many genes with relatively high fold change are predicted but not expected to change without the model. **(c)** Functional pathway analysis for predicted differentially bound genes. Enriched GO terms for predicted genes are associated with cell cycle and DNA Damage.

Upon comparison of the different classes, we observe across various TFs that differentially bound Expected/Not Predicted genes have significantly lower average change in gene expression than the Expected/Predicted genes (**Fig. 5b, SFig. 10**). Furthermore, a large number of genes in the Not Expected/Predicted category show significantly higher changes in average gene expression in comparison to all other categories, including the Expected/Predicted category for BCL3. For BCL3, the Not Expected/Predicted includes approximately 17% of the genes exhibiting differential binding and changes in gene expression that would not have been captured in the absence of the model (**Fig. 5b**).

Prior work studying the association between transcription factor binding and mRNA expression repeatedly shows a lack of change in expression in putative target genes when TF binding is disrupted.^1,2^ Our analysis reveals that a large number of these binding events might be attributed to spurious binding events and by excluding spurious binding events, we can enrich for functionally and transcriptionally relevant interactions.

To further study the biological function of these different gene groups, we conducted functional pathway enrichment analysis using Fisher’s exact test for Gene Ontology (GO) terms associated with biological processes. We identified hit GO terms as those with adjusted p<0.05. Predicted genes not only display higher number of hit terms but also had lower associated p-values for these terms (**SFig. 11**). Consider BCL3, a transcriptional co-activator of the NK-*κ*B pathway, commonly associated with apoptosis, cell proliferation and viability.^48,49^ At a stringent threshold of p<10^−5^, we find several terms associated with “cell cycle” or “DNA” (**Fig. 5c**). At this stringent threshold, none of the GO terms show significant enrichment for the expected gene set. Additionally, for predicted genes hit GO terms with p *<* 10^−15^, genes such as AKT1, MTOR and TGFB1 appear in more than 75% of terms. Prior research indicates BCL3 is often co-regulated with these genes potentially indicating a transcriptional feedback system between BCL3 and these genes that remains to be explored.^49–51^

## 5 Discussion

Biological molecules are sticky. Nucleic acids rely on only 5 different bases, and proteins only 20 different residues. While this apparent limitation induces much spurious binding, much of that binding will be short-lived. Unfortunately, many high-throughput experimental techniques will freeze those ephemeral binding events, making them visible to downstream analysis.^10,12,13,15^ The challenge to the quantitative biologist is to, after the fact, separate the spurious from the biologically relevant.

We tackled this challenge in the context of ChIP-seq data and the identification of biologically-relevant TF–target gene interactions. Errors occurring at multiple stages during ChIP-seq (**Fig. 1**) can percolate through the analysis pipeline and yield a mixture of spurious and biologically-relevant information. Here, we took a principled approach to this problem. We reasoned about the functional form of the spatial distribution of called peaks and what such functional form would tells us about how to distinguish spurious and biologically-relevant interactions.

One of the important features of our two-state model is its emphasis on the identification of dataset specific parameters. Current computational pipelines often use heuristic approaches to fixing parameter values which are then used across different cell lines or samples and different TFs across a single cell line (**Figs. 2c and S5**). This approach fails to account for variation in TF abundance across cell lines and contexts^39^ or in accessibility of regions around the target gene TSS. In contrast, our principle approach ensures that the interaction distance threshold used to distinguish spurious from biologically-relevant interactions accounts for TF and context specific variations and identifies independent parameters for each dataset

Over the last decade, several studies have attempted to link differential TF binding to concurrent changes in gene expression during molecular perturbations.^1,2^. One of the recurrent conclusions in many of these works has been that only a small fraction of binding events are responsible for regulating gene expression^3^. A large number of binding events fail to elicit a change in mRNA expression when binding is prevented through TF knockdown experiments. Here, we demonstrate that a large number of these binding events are likely to be associated with spurious binding sites. We observe that this subset can be close to 70% of the identified differential binding events and these genes display significantly lower fold change in mRNA expression than binding events deemed as biologically relevant by the model. Our model not only serves as a framework to refine functional binding events but also identifies cases that would have been overlooked using conventional methods.

While our model allows us to distinguish spurious binding events from transcriptionally relevant ones, the biological importance of variable spurious binding across different contexts remains very much an open question. For instance, are biologically-relevant binding events associated with specific chromatin contexts such as specific histone marks, DNA methylation marks or regions that are physically organized into domains in the nucleus?^47,52,53^ While we use the term “spurious” to define TF binding events that don’t directly influence transcription, these binding events can be indirectly involved in the creation and maintenance of chromatin compartments that are compatible with transcription.^22,54^ The variability of spurious binding rates across different cell lines for a given TF implies that more complex regulatory mechanisms exist that determine the binding dynamics of a TF. Furthermore, variation in spurious binding can confer evolutionary advantages or alter the ability of a cell to respond to selection pressures over large timescales.^45,46,55,56^ However, changes in spurious binding displayed by a TF can change within shorter timescales — within minutes or hours of a perturbation. How these changes influence the ability of a cell to survive in response to a chemical perturbation and whether cells with higher spurious binding are better at responding to a wider range of perturbations remains to be understood.

## 6 Code Availability

All scripts used to download data and generate the figures in this paper can be found on https://github.com/amarallab/TF_Positional_Distribution_Model.

## 7 Acknowledgments

This research was supported in part by grants from the NSF (DMS-2235451) and Simons Foundation (MPS-NITMB-00005320) to the NSF-Simons National Institute for Theory and Mathematics in Biology (NITMB). M.P. gratefully acknowledges support from the Ryan Fellowship and the International Institute for Nanotechnology at Northwestern University.

## Supplementary Figures

**Figure S1.**
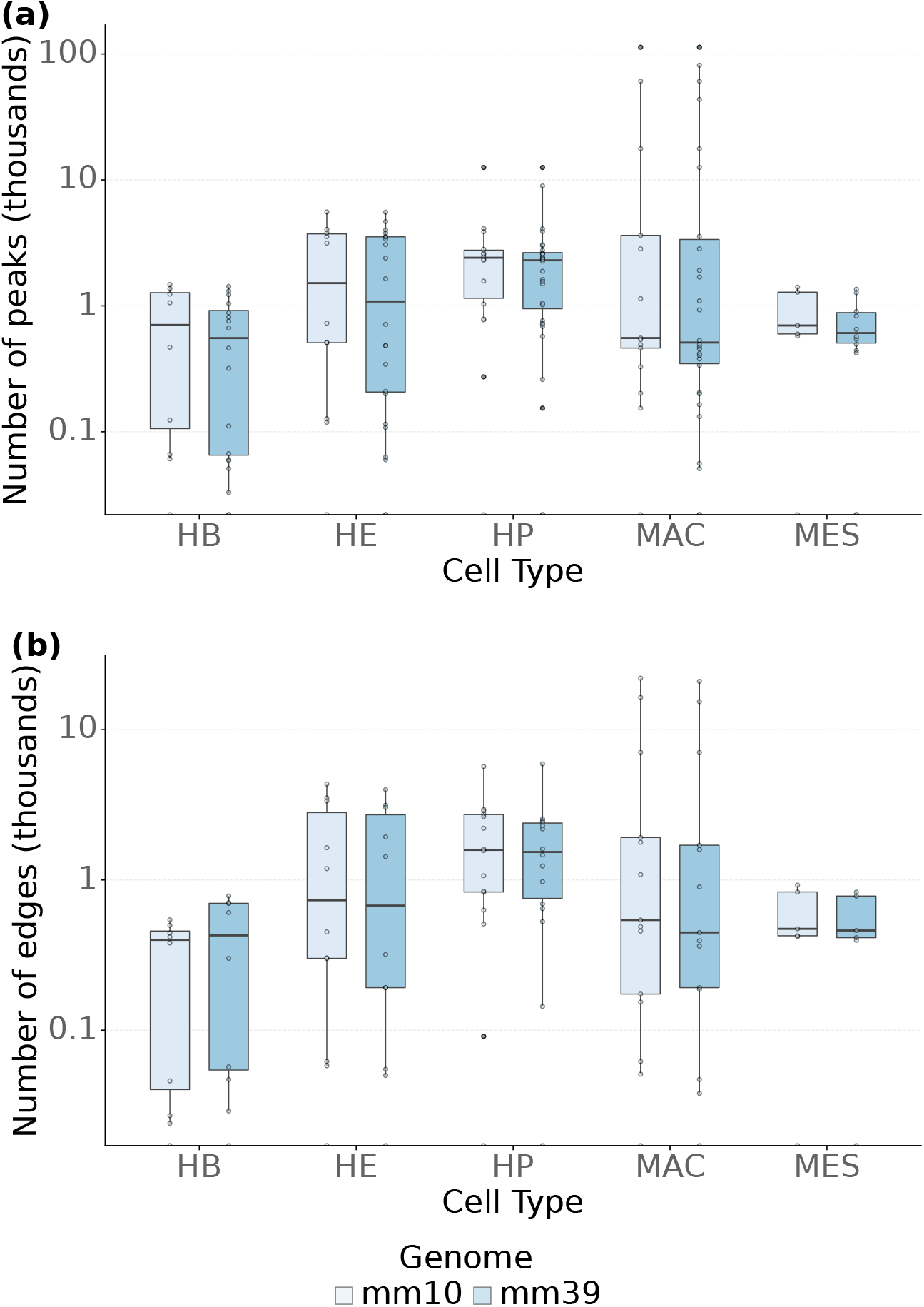
Sensitivity of called peaks to reference genome for TF ChIP-seq datasets for 5 cell types from Goode *et al*. (haematopoeitic cell differentiation). Effect of variable reference genome versions (mouse genome mm10 and mm39) on **(a)** number of called peaks and**(b)** number of edges in the TF–gene interaction network. The data suggest no significant difference arises from reference genome choice.

**Figure S2.**
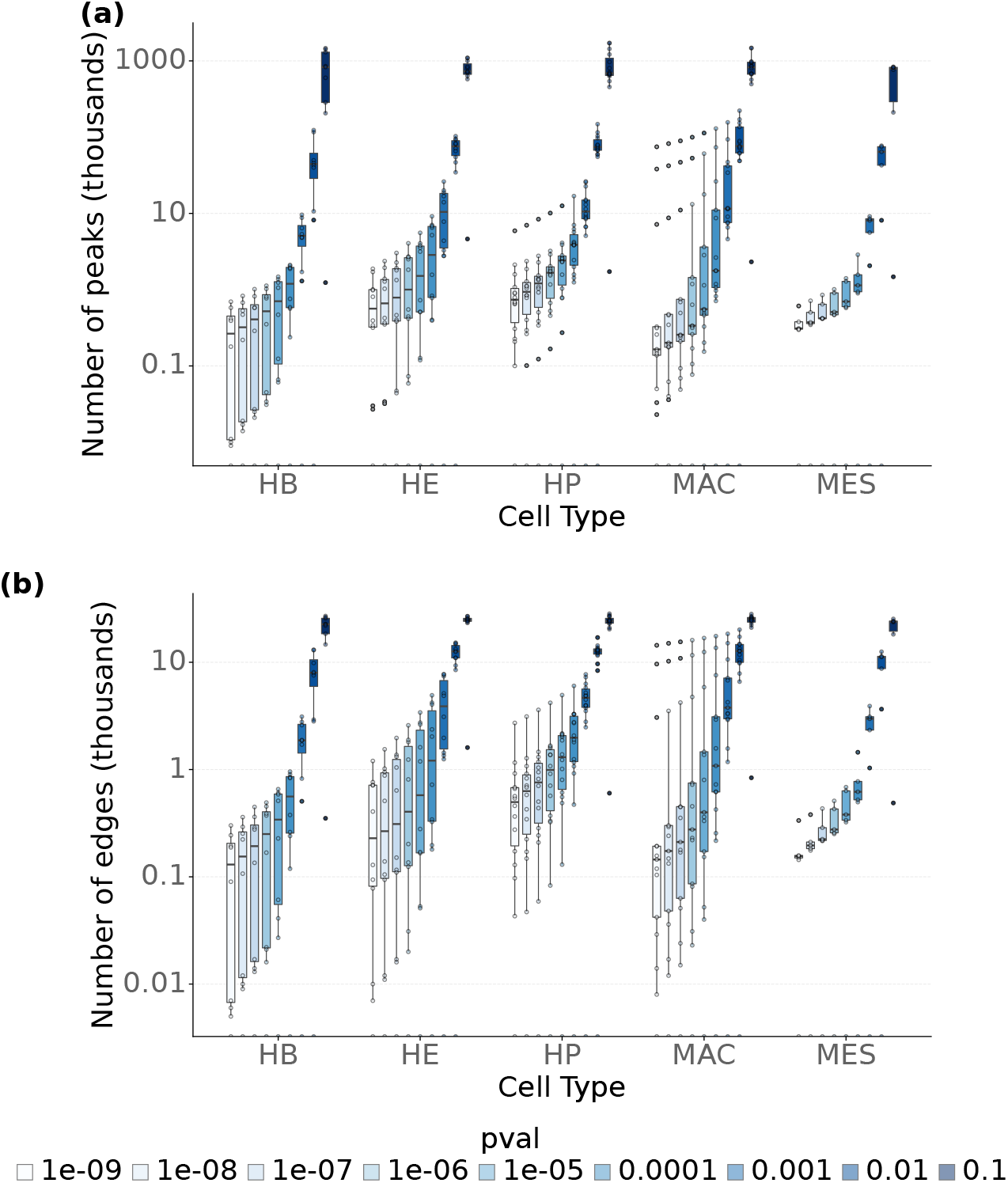
Sensitivity of called peaks to signal-to noise threshold p-value for TF ChIP-seq datasets for 5 cell types from Goode *et al*. (haematopoeitic cell differentiation). Effect of variable signal-to noise threshold p-value on **(a)** number of called peaks and **(b)** number of edges in the TF–gene interaction network. We selected p-values ranging from 10^−9^ to 10^−1^. The number of called peaks increases dramatically when *p* exceeds 10^−5^.

**Figure S3.**
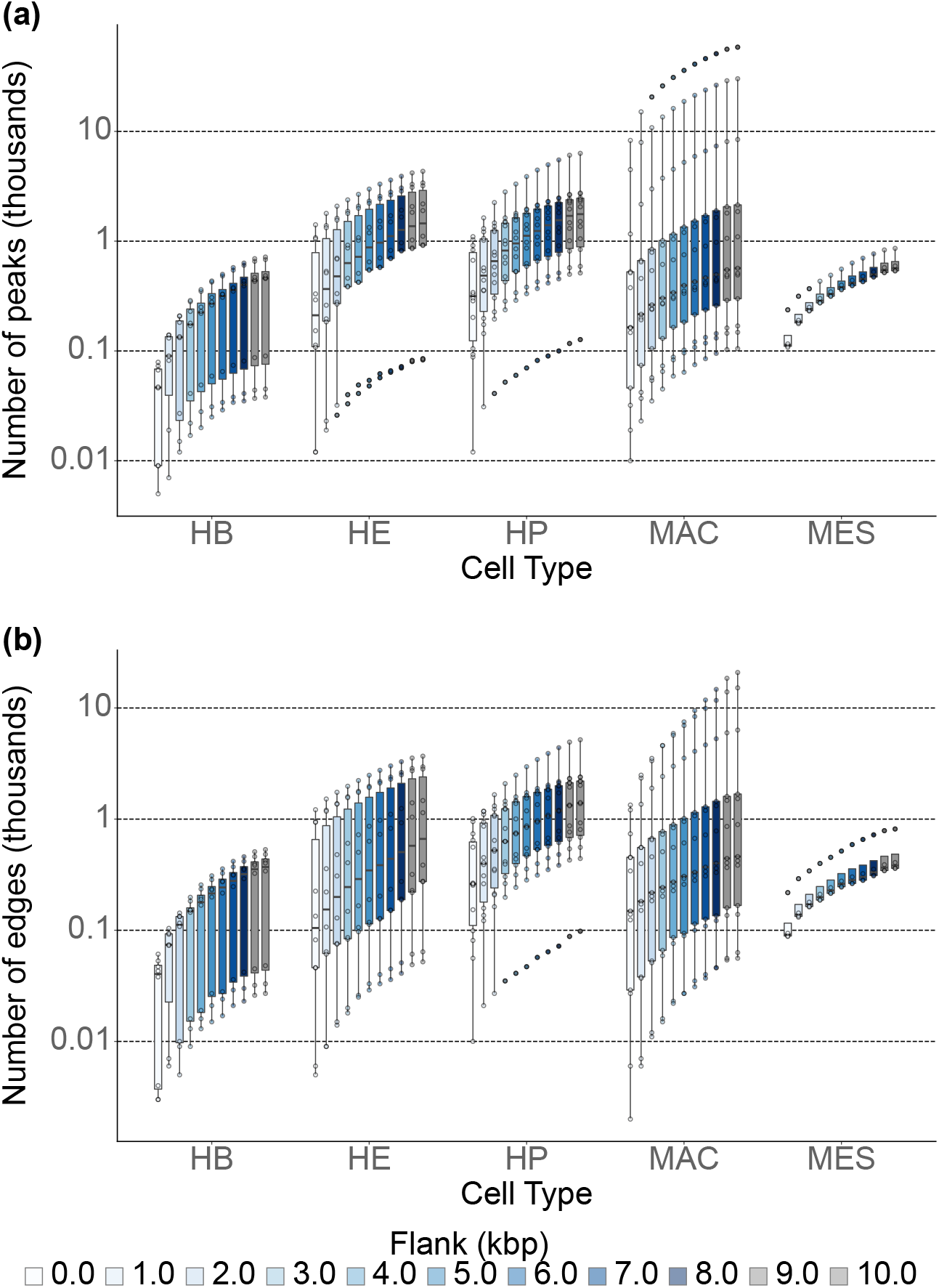
Sensitivity of called peaks to interaction distance for TF ChIP-seq datasets for 5 cell types from Goode *et al*. (haematopoeitic cell differentiation). Effect of varying interaction distance from TSS for a TF–target gene interaction on **(a)** number of called peaks and **(b)** number of edges in the TF–gene interaction network. We varied the interaction distance upto 10 kbp from the TSS in 1 kbp increments. The number of called peaks increases with the interaction distance threshold.

**Figure S4.**
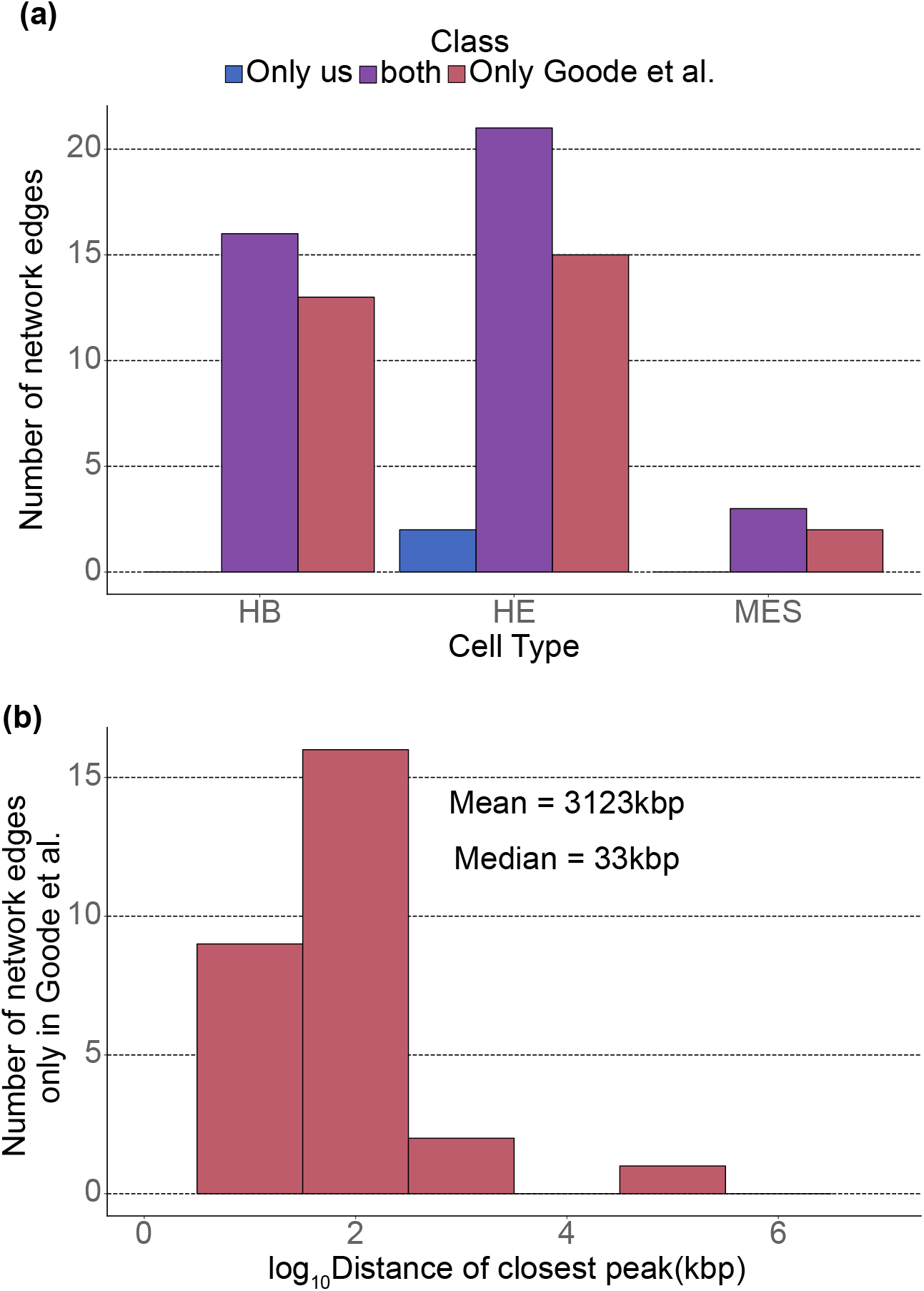
Identifying haematopoeitic cell differentiation GRN edges reported in Goode *et al*. **(a)** Number of GRN edges that were identified in Goode *et al*. only, by both analysis, and by us alone. **(b)** Distance of the nearest TF peak for the target genes those peaks identified only by in Goode *et al*. The median distance of the closest peak is 33 kbp.

**Figure S5.**
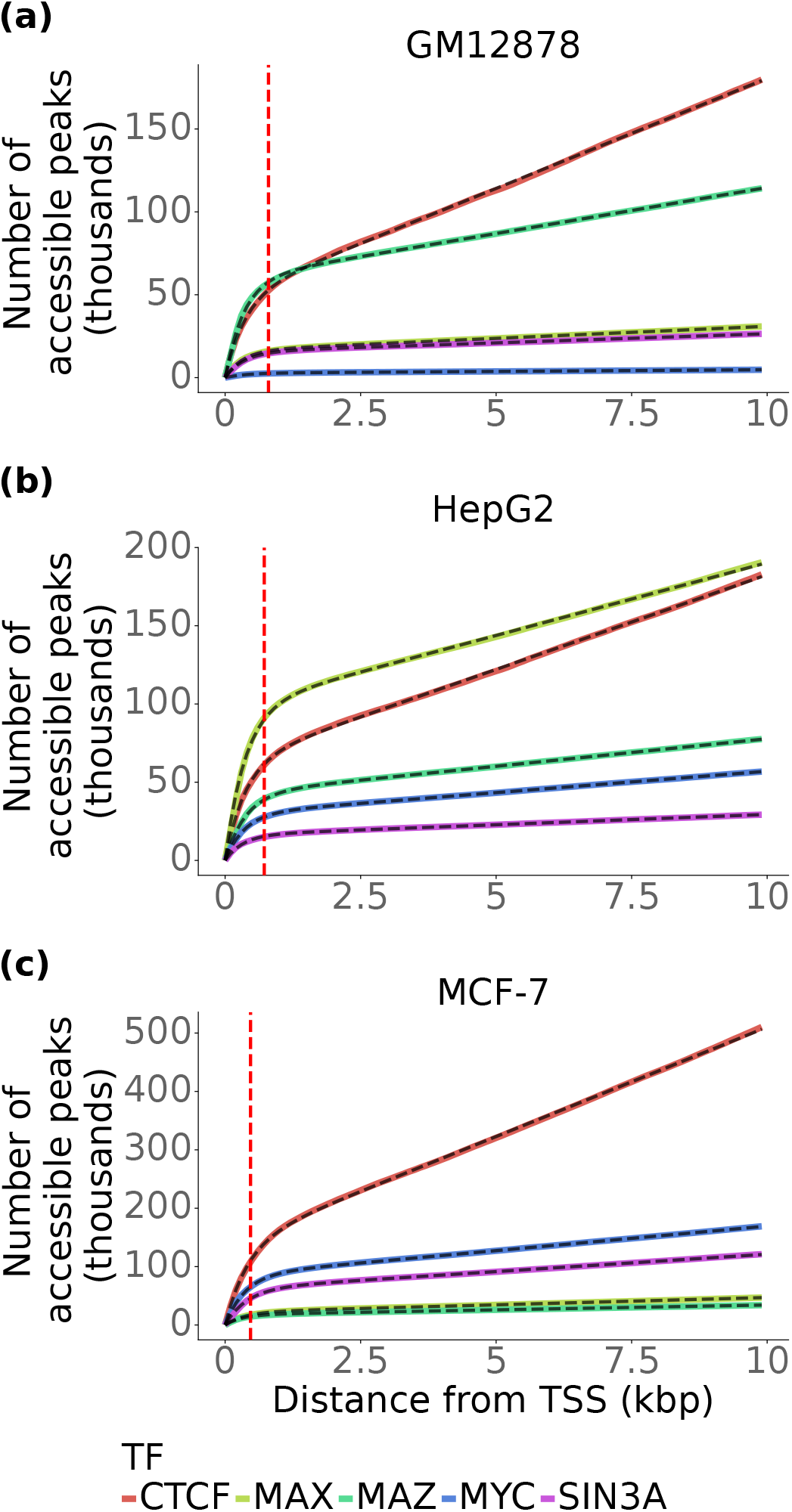
Two-state model accurately captures positional distribution of called peaks across different cell lines and TFs. Cumulative distribution of called peaks for varying distance from TSS displays two distinct phases across TFs (different colors) and different cell lines – **(a)** GM12878 **(b)** HepG2 and **(c)** MCF-7. Dashed black line represents the fit for each curve.

**Figure S6.**
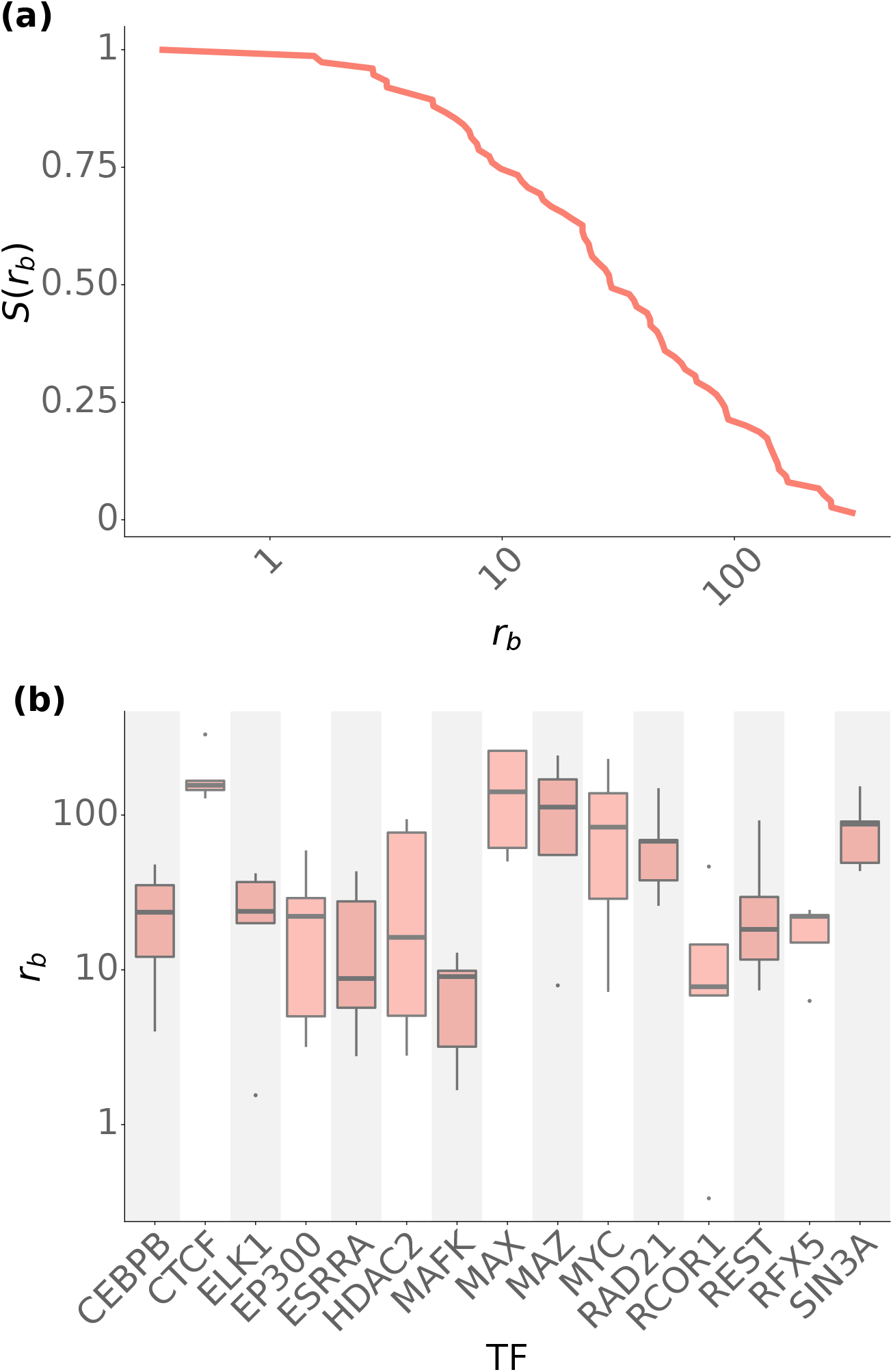
Distribution of two-state model parameter *r*_*b*_. **(a)** Survival function for *r*_*b*_ across 15 TFs in 5 cell lines. **(b)** Distribution of *r*_*b*_ within individual TFs across 5 cell lines. We observe large scale differences in values across different TFs, as well as within a TF for different cell lines.

**Figure S7.**
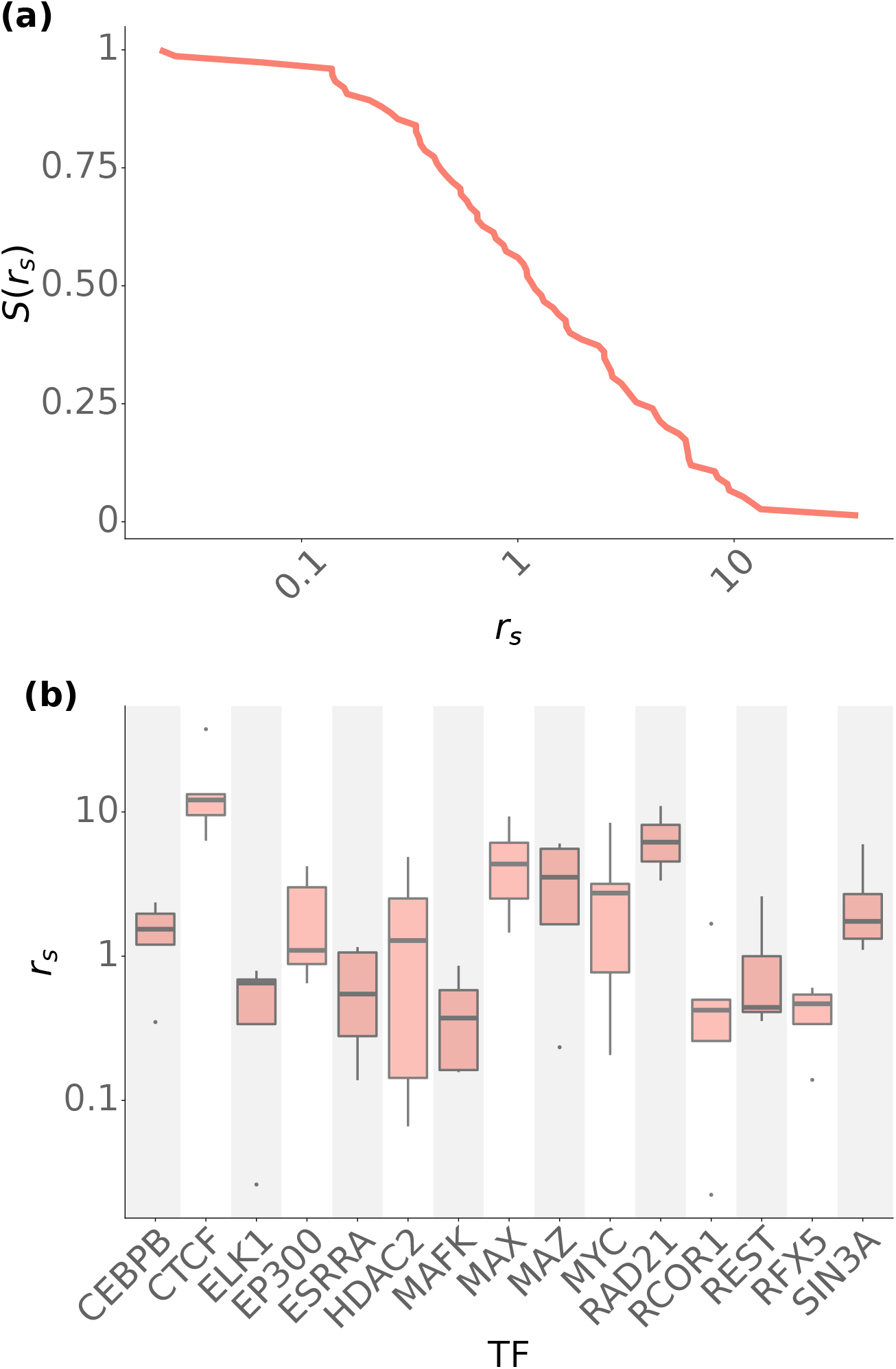
Distribution of two-state model parameter *r*_*s*_. **(a)** Survival function for *r*_*s*_ across 15 TFs in 5 cell lines. **(b)** Distribution of *r*_*s*_ within individual TFs across 5 cell lines. We observe large scale differences in values across different TFs, as well as within a TF for different cell lines.

**Figure S8.**
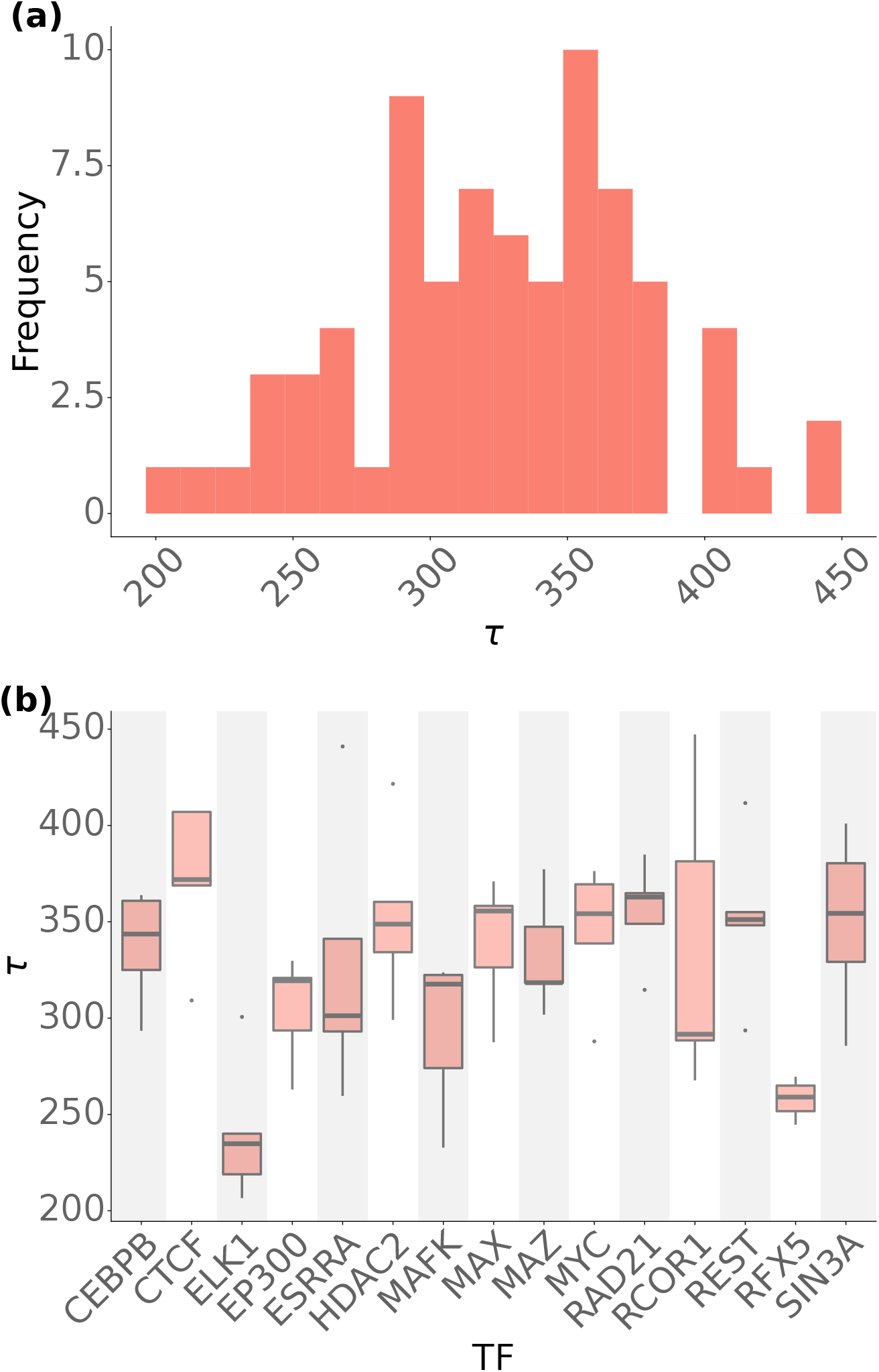
Distribution of two-state model parameter *τ*. **(a)** Distribution of *τ* across 15 TFs in 5 cell lines. **(b)** Distribution of *τ* within individual TFs across 5 cell lines. We observe large scale differences in values across different TFs but relatively smaller variations within a TF for different cell lines.

**Figure S9.**
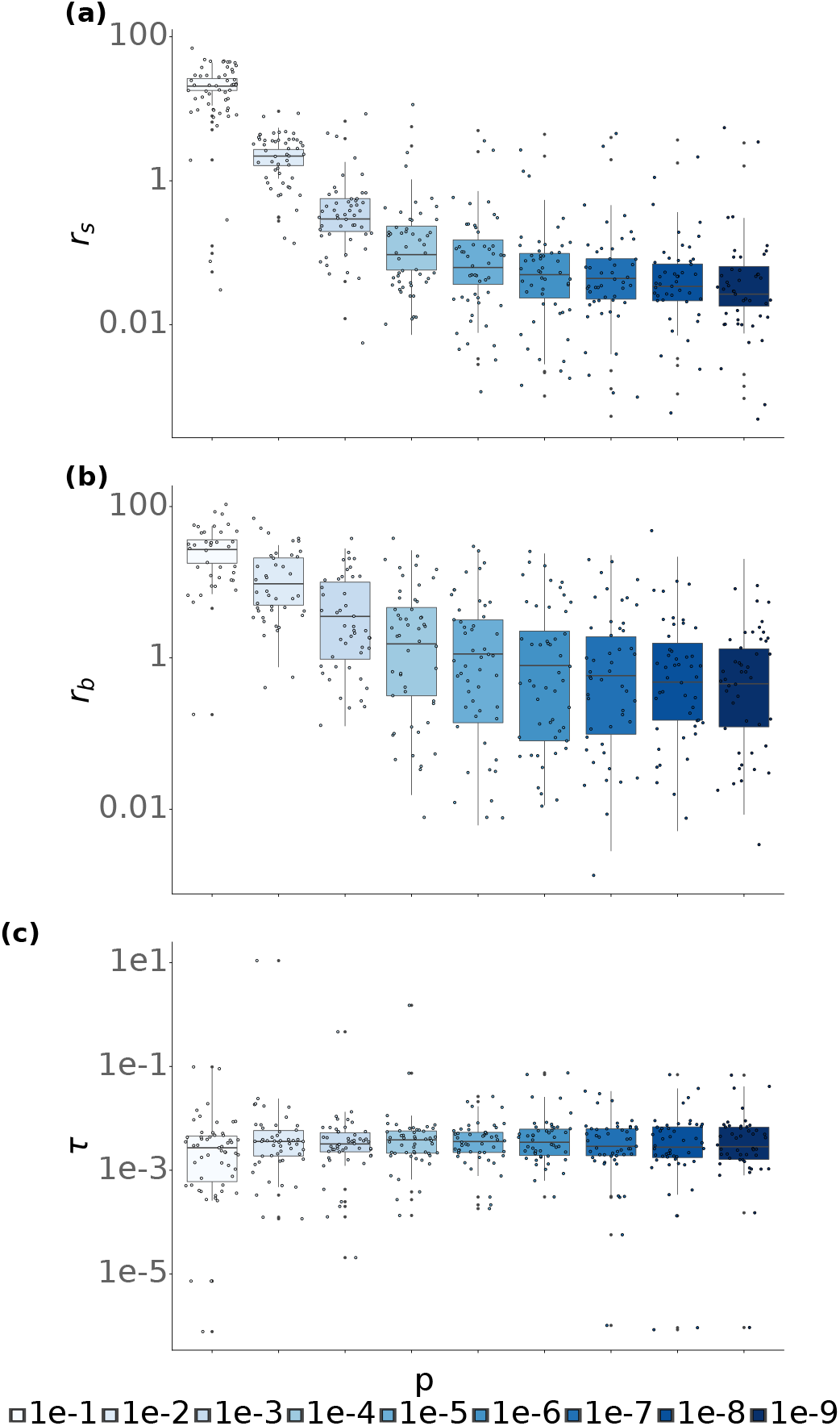
Signal-to-noise threshold p-value affects parameters associated with signal abundance but not the positional distribution. Change in **(a)** spurious binding rate (*r*_*s*_) **(b)** Scaling factor for biological binding ( *r*_*b*_) **(c)** Decay constant (*τ*) for varying p-value cut-offs for GSE69101 (haematopoeitic cell differentiation, Goode *et al*. ). While *r*_*b*_ and *r*_*s*_ display a large change with p-value cutoffs, *τ* which indicates the positional distribution around the TSS does not vary significantly.

**Figure S10.**
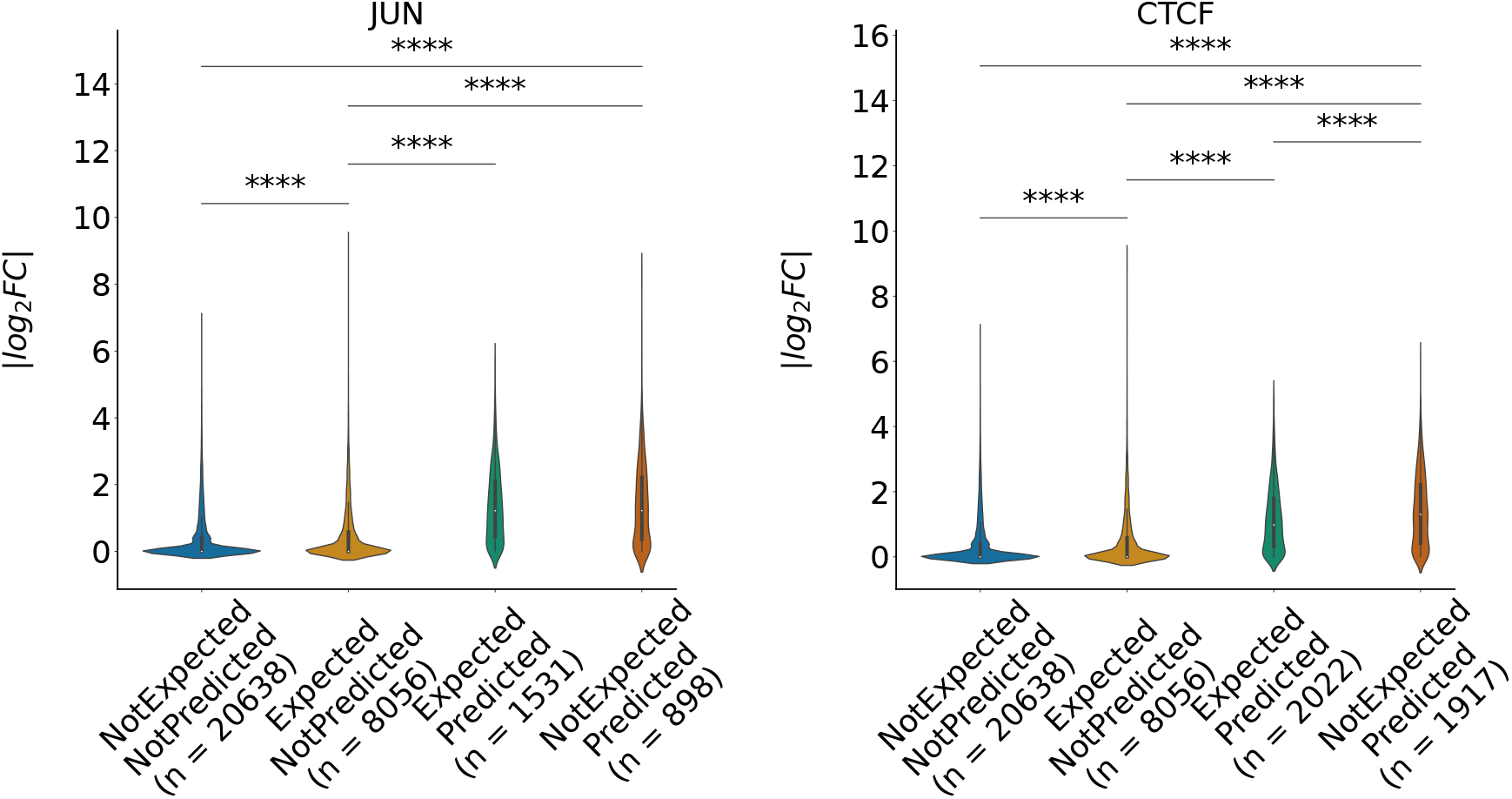
Two-state model captures differentially bound genes during 100nM dexamethasone treatment. Change in Distribution of average *log*_2_ fold change in mRNA expression between treated and untreated samples for target genes of JUN and CTCF . Many genes with relatively high fold change are predicted but not expected to change without the model.

**Figure S11.**
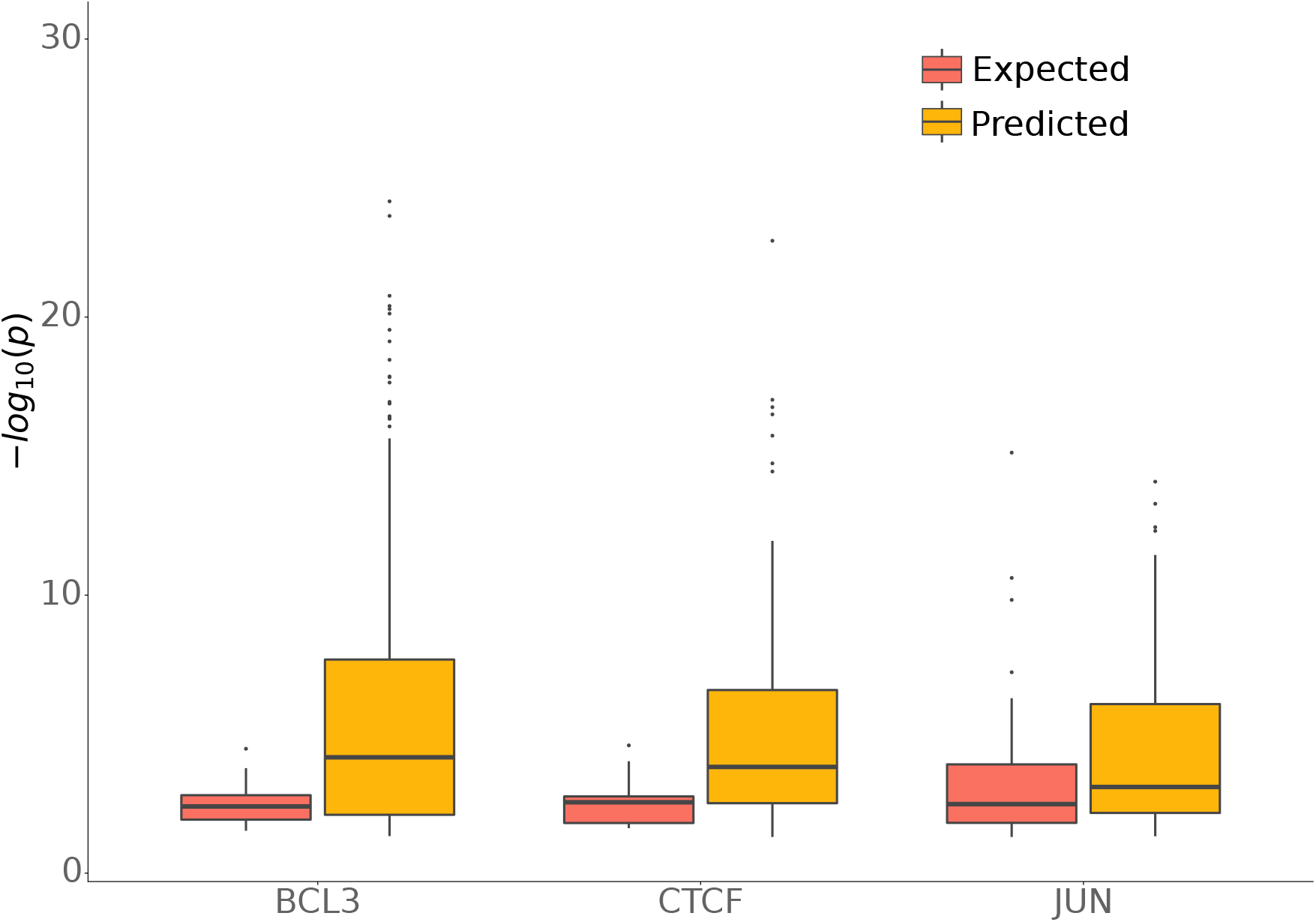
Functional pathway analysis for expected and predicted differentially bound genes. Fisher’s exact test adjusted p-value for over represent GO terms in expected and predicted differentially bound genes. Predicted genes have lower p-values associated with hit terms.

## Footnotes

1 We would like to note that spurious binding does not necessarily refer to biologically irrelevant binding events. Existing studies indicate that binding events that are not directly linked to transcription can still be crucial for other indirect functions such as chromatin organization and transcriptional condensate formation^22^

## Notes

### Competing Interest Statement

The authors have declared no competing interest.

